# ImmuneMirror: a Machine Learning-based Integrative Pipeline and Web Server for Neoantigen Prediction

**DOI:** 10.1101/2023.02.09.527828

**Authors:** Gulam Sarwar Chuwdhury, Yunshan Guo, Chi-Leung Chiang, Ka-On Lam, Ngar-Woon Kam, Zhonghua Liu, Wei Dai

## Abstract

Neoantigens are derived from tumors but are absent in normal tissues. Emerging evidence suggests that neoantigens can stimulate tumor-specific T-cell-mediated antitumor immune responses, and neoantigens are potential immunotherapy targets. We developed ImmuneMirror as a stand-alone open-source pipeline (https://github.com/weidai2/ImmuneMirror/) and a web server (http://immunemirror.hku.hk/App/) incorporating a balanced random forest model for neoantigen prediction and prioritization; the model was trained and tested using known immunogenic neopeptides collected from 19 published studies. The area under the curve (AUC) of our model was 0.87. We utilized ImmuneMirror in gastrointestinal tract cancers and discovered a subgroup of microsatellite instability-high (MSI-H) colorectal cancer (CRC) patients with a low neoantigen load but a high tumor mutation burden (TMB>10 mutations per Mbp). Although the efficacy of PD-1 blockade has been demonstrated in advanced MSI-H patients, almost half of such patients do not respond well. Our study may identify MSI-H patients who do not benefit from this treatment. Additionally, the neopeptide YMCNSSCMGV-TP53^G245V^, derived from a hotspot mutation restricted by HLA-A02, was identified as an actionable target in esophageal squamous cell carcinoma (ESCC). This is the largest study to comprehensively evaluate neoantigen prediction models using experimentally validated neopeptides. Our results demonstrate the reliability and effectiveness of ImmuneMirror for neoantigen prediction.

## INTRODUCTION

Immunotherapy uses the immune system to detect and fight against cancer cells. Accumulating evidence shows that the presence of neoantigens derived from somatic mutations in tumor cells elicits a potent immune response as a part of antitumor immunity through specific cytotoxic T cells (1,2). Previously, various methods have been proposed for neoantigen identification, such as MHCflurry (3), NetMHCpan (4–6) and NN-Align (7), which predict the binding affinity between peptides and their corresponding major histocompatibility complex (MHC) alleles. Binding affinity is a good reference to prioritize neoantigens because MHC classes I and II help the immune system bring the bonded complex to the surface of cancerous cells for recognition by T cells. Therefore, binding to MHC molecules is a prerequisite for immunogenicity. However, the actual variant expression, HLA presentation, peptide processing, and transportation, as well as the ultimate T-cell response to these neoantigens, have not been considered in these existing binding affinity-based tools, therefore, these previous methods may fail to provide reliable predictions in real-world scenarios. Recently, by integrating peptide features, Wells *et al*. developed a model of tumor epitope immunogenicity to filter out nonimmunogenic peptides, and the results improved the effectiveness of neoantigen prediction (8). This model is based on stringent cutoffs for several selected features, including binding affinity, binding stability, tumor abundance, the ratio of binding affinity between mutant and wild-type peptides (9), and T-cell receptor recognition probability (foreignness); the model showed promising results with precision (true positive/(true positive + false positive)) above 0.7. However, Wells’ study (8) used different criteria during the training and validation steps to filter neoantigens, making it difficult to implement with other data sets. A continuous effort is still needed to further improve the prediction accuracy for clinical application by incorporating more relevant biological features that are involved in these complicated biological processes.

In this study, we developed ImmuneMirror, an all-in-one multiomics data analysis bioinformatics pipeline, to access the key genomic and transcriptomic features associated with the cancer immunotherapy response. The ImmuneMirror pipeline and web server 1.0 incorporate a machine learning model to incorporate significant biological features for neoantigen prediction. With this advanced machine learning model trained by known neopeptides with T-cell immunogenicity, ImmuneMirror overcomes the issue of unbalanced neoantigen distribution, i.e., immunogenic mutation-derived neoantigens are relatively rare compared to the total number of mutations detected. We applied ImmuneMirror to real-world data to systematically investigate neoantigens in gastrointestinal tract (GIT) cancers using matched whole-exome sequencing (WES) and RNA Sequencing (RNASeq) data; furthermore, we compared the results with putative neoantigens that are derived from hotspot mutations in cancer-related genes restricted by the following four common HLA alleles: HLA-A02:07, HLA-A24:02, HLA-A02:01, and HLA-A11:01. The top candidate neopeptide, YMCNSSCMGV-TP53^G245V^, derived from a hotspot mutation restricted by HLA-A02, was evaluated experimentally.

## MATERIALS AND METHODS

### Selection of machine learning models for neoantigen prediction

To build the prediction model for identifying neoantigens and incorporating more relevant genomic and transcriptomic features, we first gathered a list of neopeptides with experimentally confirmed T-cell responses to use as the training data for model construction.

The binding affinities of peptide amino acids were predicted through pVACtools (10) with multiple prediction methods for MHC class I. Feature selection was performed based on hypothesis tests to include the neoantigens that were detected by pVACtools (10); the hypothesis tests were the median binding affinity score of the epitope estimated from all prediction algorithms as well as the mean hydropathy of the last 7 residues on the C-terminus of the peptide. Additionally, considering that binding affinity is not the only parameter governing tumor epitope immunogenicity, we added the following relevant features for thorough analysis and to improve prediction accuracy: ‘agretopicity’ (9,11) and ‘foreignness’ (12–14), hydrophobicity, binding stability, peptide processing, and transportation scores. The final training data set included a total of 1199 peptides that were tested *in vitro*, 93 of which had positive T-cell responses. Ten of the 211 tested peptides were immunogenic. These neopeptides were identified from 19 published studies (**Supplementary Tables S1 and S2**).

The class distribution of the data set is extremely unbalanced due to the low proportion of immunogenic neoantigens, which activate T cells. Consequently, conventional classification algorithms are largely affected by the majority class and thus give biased attention to the minority class, resulting in poor prediction performance. Therefore, we adapted balanced random forest learning algorithms to address this issue (15). These methods were evaluated using the area under the receiver operating characteristic curve (AUC) metric.

Random forests are ensemble learning algorithms for classification that construct multiple decision trees during the training process. Given a training set Z = z_1_, …, z_*n*_, where *Z*_i_ = (*x*_i_, *y*_i_) ∈ *X* × *Y*, the algorithm repeatedly (*B* times) selects a bootstrap random sample with replacement and a random subset of the predictive features from the training data set and then fits decision trees *f*_b_ to each of those bootstrap random samples *X*_b_, *Y*_b_,. When building decision trees, a random number of *m*predictors are selected as split candidates from the entire *p* predictor pool; typically, we set 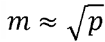. The Gini index, defined as 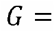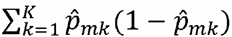, where 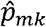 is the proportion of training observations in the *m*th region belonging to the *k*th class; the Gini index is used as a criterion to make the binary split when growing a tree. The final classification output is based on majority voting from all the base decision trees or the classification can be made by taking the average of all predictions using the following equation: 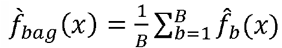 (16)

### SMOTE and under-sampling combination + random forest

Conventional random forest uses the standard bootstrap sampling strategy with equal sample probability for each observation, and this strategy does not perform well for an extremely imbalanced data set. Therefore, we proposed modifying the random forest algorithm using the advanced resampling technique, which over-samples from the minority class and under-samples observations from the majority class to increase the minority-majority ratio from approximately 1:12 to 1:3. This under-sampling step can be achieved using the *smote_and_undersample* function in the R package *hyperSMURF* (17). This function first generates synthetic examples based on the synthetic minority over-sampling technique (SMOTE), which retains each minority class sample and introduces synthetic examples along the line segments joining some of the *k* minority class nearest neighbors (18). In our case, we set the multiplicative factor *f_p_* to 2 and *k* to 5, so two neighbors from the five nearest neighbors were selected. Then, observations from the majority class were under-sampled to reach the preset class ratio. We then fit the conventional random forest on the resampled data. These processes were repeated several times to guarantee that most of the data were involved in the training process. Finally, we selected the model with the best AUC (0.8294) using the testing data (**Figure 1**).

**Figure 1.**
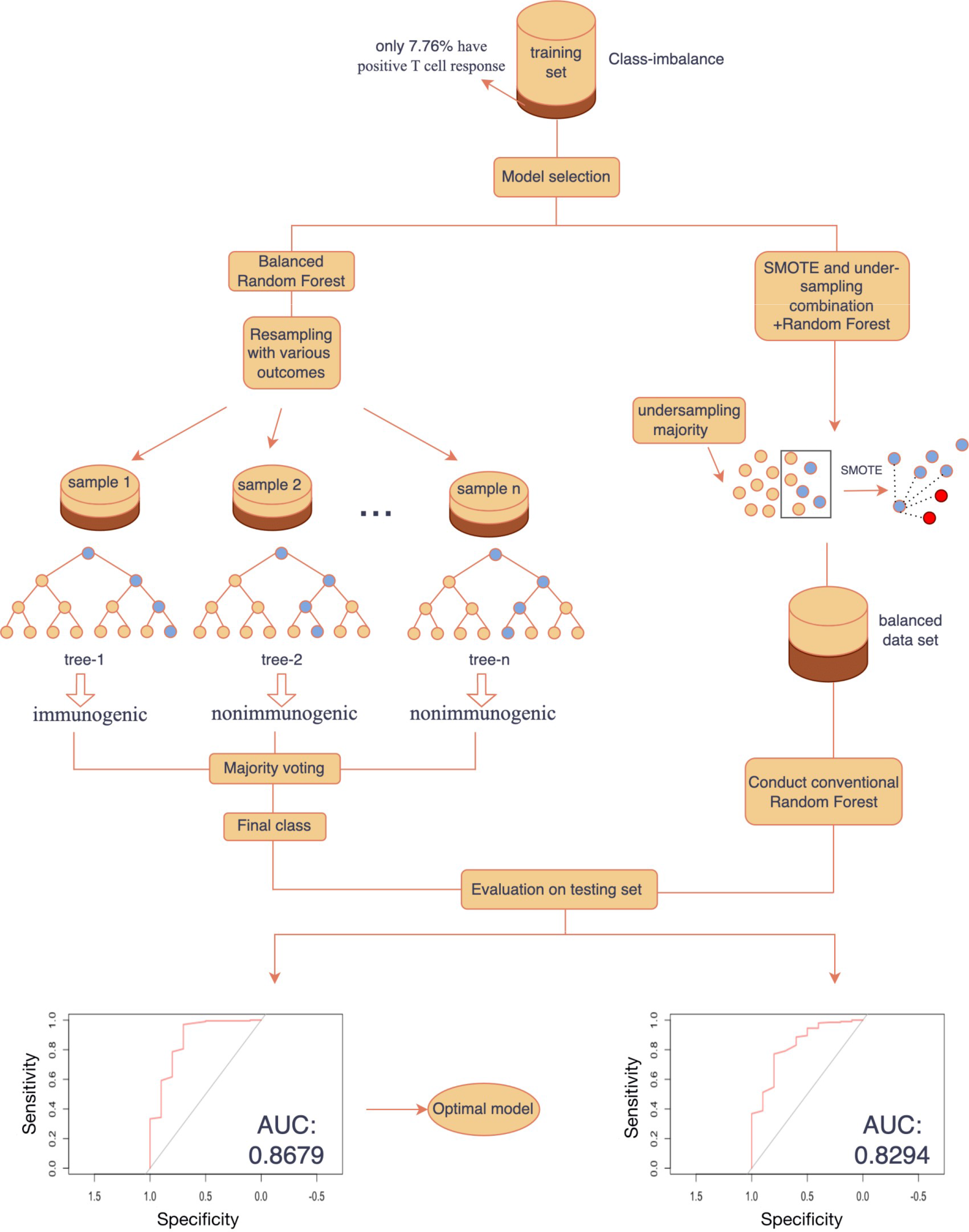
Evaluation of different random forest algorithms used for neoantigen prediction with AUC performance: Left panel: Balanced random forest; Right panel: Random forest with under sampling.

### Balanced random forest

To address the imbalanced data issue, Chen and Breiman (19) developed the balanced random forest algorithm, which substantially improved the performance of the random forest algorithm by replacing the equal-weight sampling strategy with random under-sampling for decision tree formation. More specifically, for each iteration in a random forest, we randomly drew a bootstrap sample from the minority class and obtained the same number of cases from the majority class with replacement. Then, we formed a classification tree based on the balanced data. We repeated the above steps many times and then determined the final prediction via majority voting (19). The balanced random forest algorithm was implemented in the *train* function in the R package *caret* (20), and the optimal value of parameters was tuned by a 5-fold cross-validation method. The AUC on the testing set was 0.8679 (**Figure 1**). By evaluating the above two models, the balanced random forest model outperformed the others, making it the optimal model to predict neoantigens.

### Peptide synthesis and quality control

The selected peptides were synthesized. The CI resins were selected and deprotected in 20% piperidine dimethylformamide (DMF) solution. The resin was filtered off and rinsed with DMF three times to remove Fmoc residues. The completeness of amino deprotection was measured by taking a sample of the resin and mixing it with detection reagents A and B. If there was a color change, the Fmoc groups were removed successfully. The amino acid solution was added into a mixture of the resin and di-isopropyl carbodiimide (DIC) in DMF, and the mixture was shaken at room temperature. The completeness of the coupling reaction was confirmed by taking a sample of the resin and mixing with detection reagents A and B, followed by resin washing with DMF three times. When all the amino acids were coupled onto the resins, the peptide chain was dissociated from the resins by treatment with TFA/DMF. The crude peptides were further purified by reversed-phase high-performance liquid chromatography (HPLC) and were frozen and dried under vacuum. The molecular weights of the selected peptides were analyzed by LC◻JMS. Endotoxin levels were detected using Horseshoe Crab Reagent. The peptide with an endotoxin level <10 EU/mg was used for the MHC binding assays.

### HLA-A02:01 peptide-binding assay

The QuickSwitchTM Quant HLA-A02:01 Tetramer Kit-PE was used to investigate the binding affinity of the selected neoepitope to MHC HLA-A02:01. The synthesized peptides were incubated with the MHC HLA-A02:01 complex, which already contained a control peptide. The tested peptide competed with the control peptide, and the exchange rate was used to identify the peptide-binding affinity. QuickSwitch™ Quant HLA-A02:01 Tetramer Kit-PE and flow cytometry were used to investigate the exchange rate of the MHC HLA-A02:01 control peptide. The tested peptide was mixed with the tetramer and peptide exchange factor for 4.5 hours at room temperature. The peptide exchange rate was quantitated by flow cytometry (21,22). The reference positive peptide was provided by the QuickSwitch™ Quant HLA-A*02:01 Tetramer Kit-PE.

## RESULTS

### Overview

The overall workflow of this study is depicted in **Figure 2**. The machine learning (ML) model was developed using the balanced random forest algorithm for neoantigen prediction with the incorporation of multiple biological features relevant to neoantigen biogenesis, transportation, presentation, and T-cell recognition (agretopicity, foreignness, hydrophobicity, binding stability, peptide processing, and transportation scores). This ML model was incorporated into the ImmuneMirror bioinformatics pipeline, which is also a web server for neoantigen prediction and prioritization from multiomics sequencing data. The pipeline takes the raw FASTQ reads as input, while the web server takes VCFs file containing the somatic mutations (**Figure 3**). This pipeline was applied to identify neoantigens derived from somatic mutations in cancer-related genes with common MHC class I subtypes in Pan-Cancer studies and from real-world WES and RNASeq data from GIT cancer patients. Experiments were carried out to confirm the binding affinity of the putative neoantigens with MHC class I HLA-A02:01.

**Figure 2.**
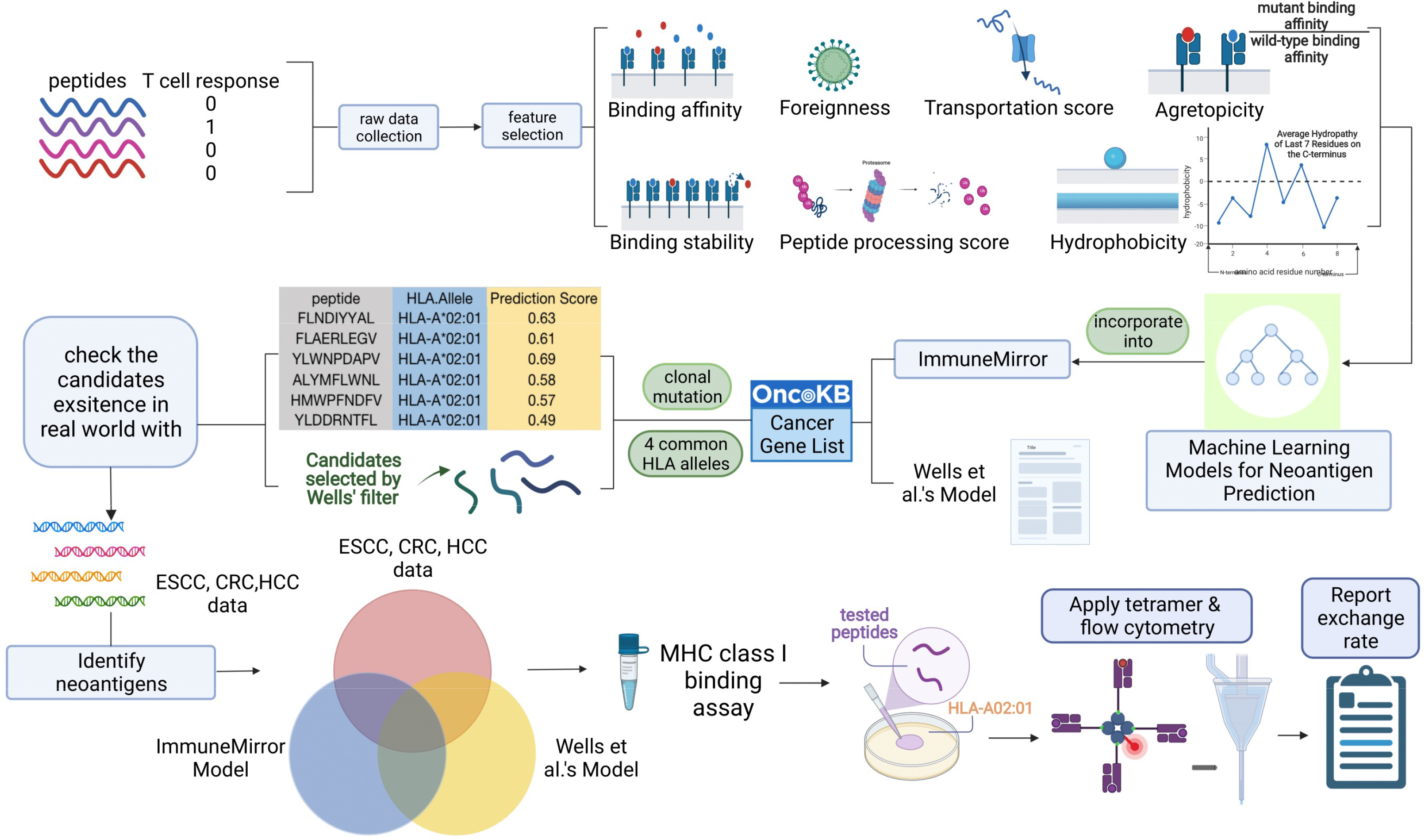
Overall study workflow. Neopeptides with experimentally confirmed T-cell responses were gathered as training data for model construction. Relevant features were selected through feature selection. The prediction model (ImmuneMirror) was established with an advanced machine learning algorithm. ImmuneMirror was subsequently applied to the hotspot mutations derived from the common cancer gene list from OncoKB to predict potential neoantigens. Wells’ criteria (8) were also applied to hotspot mutations for the selection of neoantigens. The publicly available data from ESCC, CRC & HCC patients were processed and analyzed by ImmuneMirror. We compared the results obtained from the two data resources and identified overlapping candidates that were then subject to experimental validation of binding affinity with HLA-A02.

**Figure 3.**
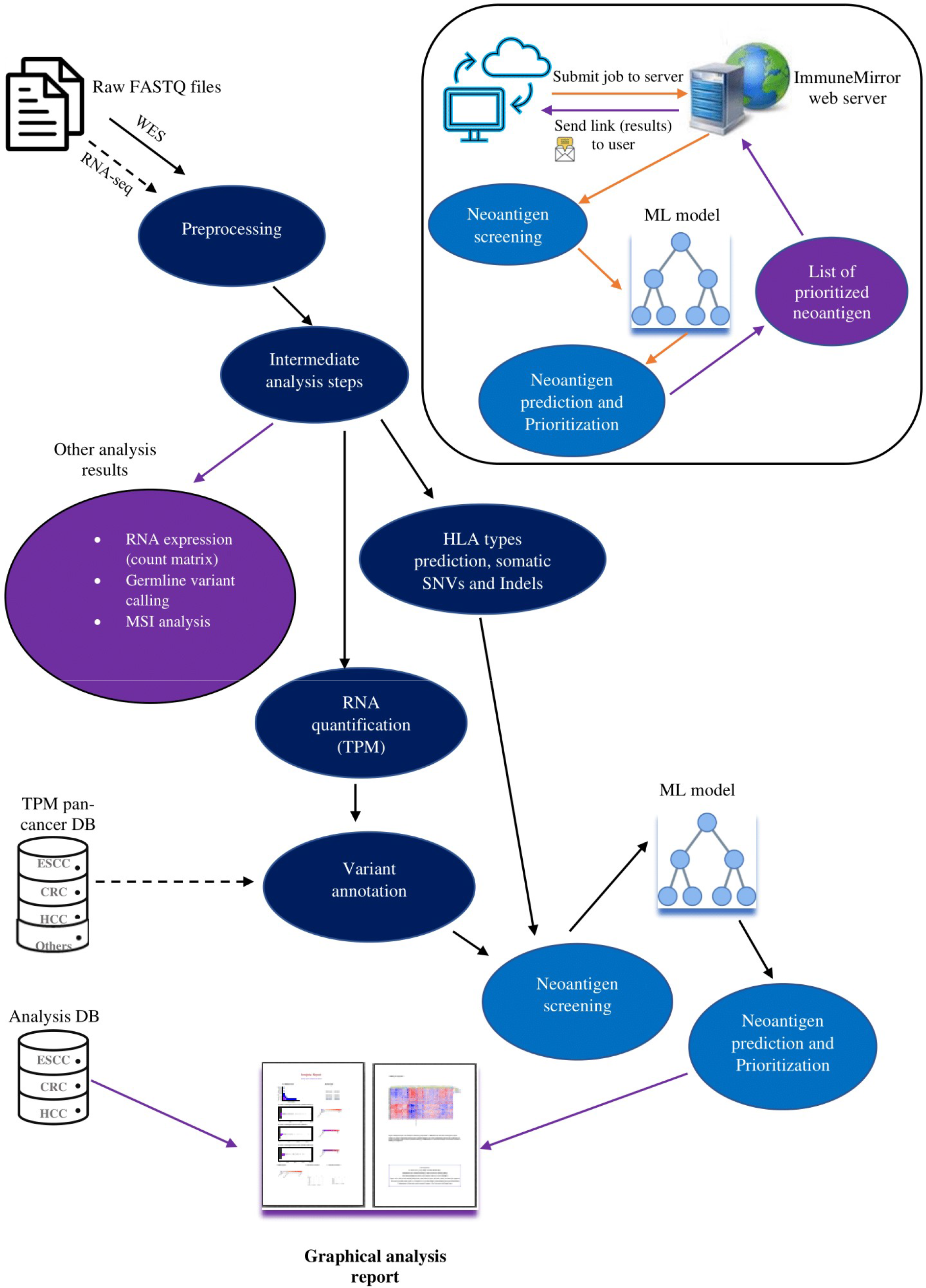
The overall workflow of ImmuneMirror, major analysis steps involved in the ImmuneMirror pipeline, and the ImmuneMirror web server. The ImmuneMirror pipeline preprocesses raw FASTQ files, including multiple analysis steps (e.g., prediction of HLA subtypes, SNV and indel detection, variant annotation, neoantigen prediction and prioritization), and generates a graphical analysis report for each sample. The input for the web server is a VCF file, and the analysis result (list of prioritized neoantigens) is sent as a web link to the email address of the end user.

### Implementation of the ImmuneMirror pipeline and web server

We developed the ImmuneMirror pipeline for neoantigen prediction and prioritization based on multiple genomic and transcriptomic features. The workflow of the ImmuneMirror pipeline is depicted in **Supplementary Figure S1**. The pipeline was built as a docker container that can be run in any docker-supported operating system, such as Linux, Mac and Windows. The pipeline required FASTQ input of matched normal-tumor WES samples and tumor bulk RNASeq samples. It was implemented with various benchmark bioinformatics packages (see Supplementary Data file for details description), such as BWA-mem (23) for WES alignment, STAR (24) for RNAseq alignment, Genome Analysis Toolkit (GATK4) (25), Picard Toolkit (http://broadinstitute.github.io/picard/), etc. for preprocessing; MSIsensor-pro (26) for microsatellite instability analysis; GATK4 (25) for germline variant calling and somatic SNV and indel detection; OptiType (27) for HLA class I haplotyping and PHLAT (28) for HLA class II haplotyping; and pVACtools (29) for predicting the binding affinity of HLA class I and II. The full list of packages and software that were used for ImmuneMirror pipeline development are listed in **Supplementary Table S3**. Finally, we implemented the prediction model as an R function for prioritizing the neoantigens restricted by HLA class I. The germline and somatic mutations, estimated tumor mutation burden (TMB), microsatellite instability (MSI) status, HLA typing, neoantigen load for HLA class I and II, the top-ranked neoantigens with T-cell immunogenicity, and the expression of the selected gene signatures are the final outputs of the pipeline. Moreover, we evaluated the accuracy of MSI status prediction by ImmuneMirror using the known MSI status generated by The Cancer Genome Atlas (TCGA) studies; the sensitivity, specificity and accuracy were calculated to be 97.37%, 99.56% and 99.25%, respectively. Users can download the Docker image and the relevant files (reference files and example samples) from http://immunemirror.hku.hk/ and clone the ImmuneMirror pipeline from the GitHub repository (https://github.com/weidai2/ImmuneMirror/).

Apart from the development of the stand-alone pipeline, we also developed an ImmuneMirror web server (**Figure 3**) that takes a VCF file containing the somatic mutations detected by MuTect2 as the input and identifies the potential neoantigens derived from somatic mutations for both HLA class I and class II molecules. Users can upload a VCF file, enter a set of alleles for both HLA class I and II, and select peptide lengths via the web interface. The uniform resource locator (URL) link for downloading the results will be sent to the user-provided e-mail automatically by the server upon job completion. The web server is freely available for users with detailed usage instructions at http://immunemirror.hku.hk/App/.

### Graphical analysis report

With the advantages of our developed analysis database, ImmuneMirror produces a visual analysis report for each of the samples. The report, as illustrated in **Supplementary Figure S2**, includes TMB, HLA types, neoantigen load for HLA class I and II, MMR status, germline and somatic mutations, and innate anti-PD1 resistance (IPRES) gene expression signature (30). The TMB is shown as the number of mutations per Mb. The HLA typing of the sample is presented in a table for class I and class II. The number of neoantigens restricted by HLA class I and class II are illustrated as box plots and bar plots with indicators for high and low neoantigen loads, respectively. The MMR status of the sample is reported. The cutoff for the MSI-high group was determined by the optimal value of the MSIsensor-pro score for distinguishing the MSI group from the other groups in CRC. The sample with a MSIsensor-pro score higher than the cutoff was defined as MMR deficient. Moreover, both germline variants and somatic mutations of selected genes, such as *BRCA2*, *B2M*, *MLH1* and *MSH2*, and expression of the genes from the IPRES signature are included in the analysis report. It has been reported that these selected mutations and gene expression signatures are relevant to the immunotherapy response (30,31).

### Testing, computation speed evaluation, resources and comparison of features

We tested the pipeline on Linux operating systems (ubuntu 20.04). It took approximately 30 hours to process one pair of samples with 13 threads. Moreover, we ran the pipeline with multiple pairs of samples from different cancer types, i.e., ESCC, HCC, and CRC. Users can run ImmuneMirror with a list of samples, and the actual run time depends on the computation speed and resources of their own devices. In general, we recommend a device with at least 64 GB of RAM and the necessary space for the pipeline, including docker image (79.6 GB), supporting files (483 GB) and analysis results (approximately 41 GB for one pair of samples), to successfully run the pipeline. The supporting files provide the necessary resources, such as the reference human genome (hg38), to run the pipeline; thus, no additional step is needed to download these files or to reconfigure the pipeline. The web server has been tested on Linux, macOS, and Windows platforms with various web browsers (**Supplementary Table S4**). The format of the input/output files and detailed instructions are provided on the website and will be updated regularly.

We compared the bioinformatics tools available for neoantigen prediction (**Supplementary Table S5**). Compared to other existing pipelines, only ImmuneMirror has all of the following six unique features: methods for prioritization, docker image, web server, neoantigen prediction for HLA class I and II, and multiple prediction algorithms. As a docker image, the ImmuneMirror pipeline takes the raw FASTQ files from both WES (matched normal-tumor pairs) and RNASeq (tumor, optional) data as the input. On the one hand, similar to pVAC-Seq (10), ImmuneMirror can be used for neoantigen prediction restricted by HLA class I and class II using multiple algorithms. On the other hand, ImmuneMirror provides a unique web server taking the VCF file that contains the somatic mutations detected by MuTect2 as the input for neoantigen prediction, which makes ImmuneMirror user friendly.

### Application of ImmuneMirror to real-world data

#### Identification of clonal mutations as neoantigens from TCGA Pan-Cancer studies

The top 27 cancer-relevant genes with hotspot mutations (frequency >0.1%) from the OncoKB (32) cancer gene list were selected for analysis. Each mutant is paired with four common HLA alleles, HLA-A02:07, HLA-A24:02, HLA-A02:01, and HLA-A11:01, derived from Asian populations. The prediction scores of the mutant-HLA combinations as well as many other biological features were calculated by ImmuneMirror. Wells *et al*. (8) developed selection criteria (binding affinity < 34 nM; binding stability > 1.4 hours; tumor abundance > 33 TPM; agretopicity < 0.1 or foreignness >10^−16^) to select neoantigens based on several experimental validation results (8). To present a thorough analysis, we also applied Wells’ criteria (8) to all the mutants identified from these 27 genes to compare with our prediction results. Neoantigen candidates were finalized if 1) the prediction score was greater than 0.515 (sensitivity: 0.851; specificity: 0.7, evaluated by the testing data set) and 2) they fulfilled Well’s criteria (8) with an adapted gene expression cutoff of TPM >10. We finally identified a total of 9 neoantigens derived from the mutations of 27 genes with a mutation frequency >0.1% from TCGA Pan-Cancer studies. The results included multiple potential neoantigens derived from TP53^P250L^, TP53C^242F^, and TP53^S241F^ (**Table 1**).

**Table 1:**
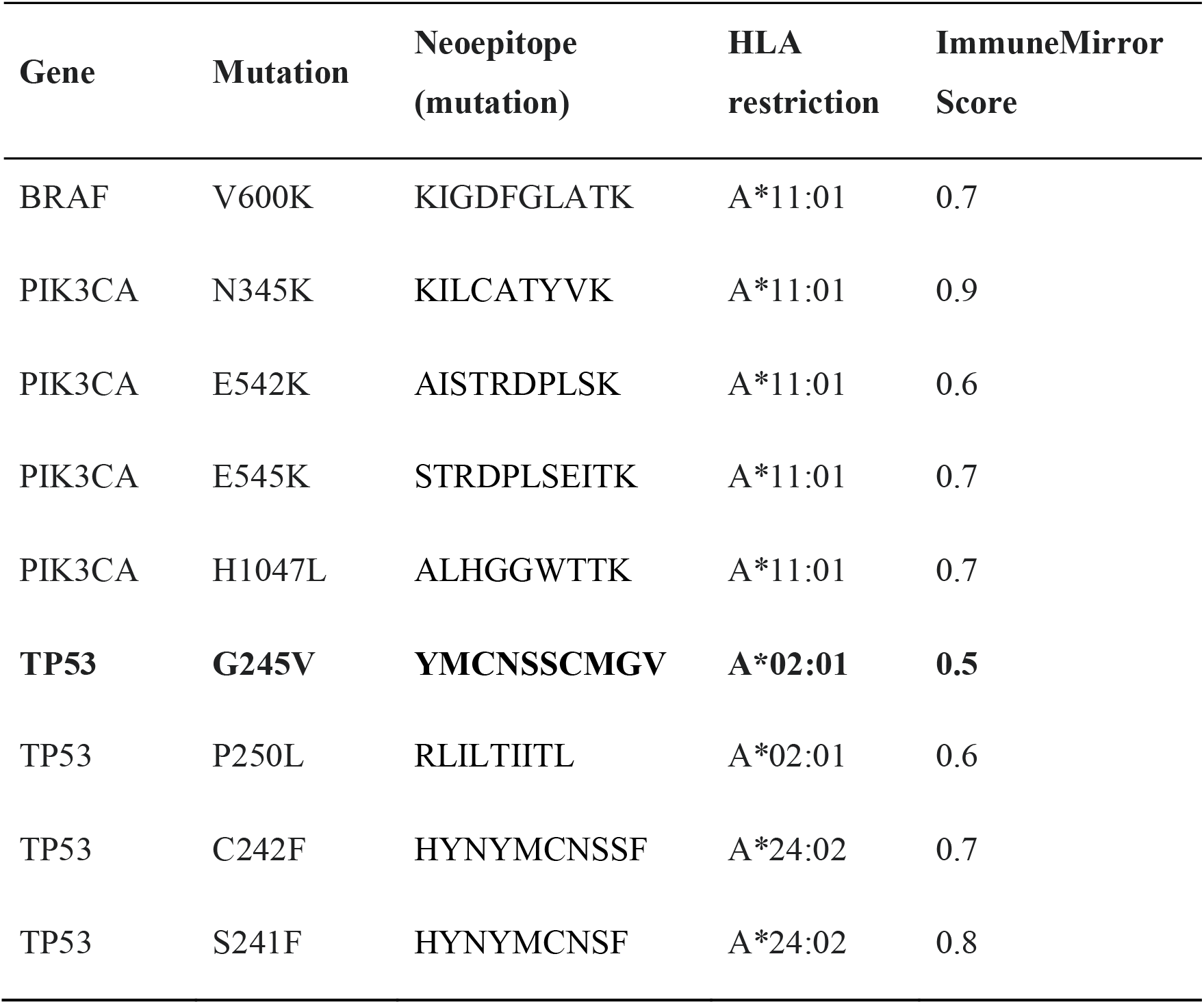
The neoantigens identified from hotspot mutations in TCGA Pan-Cancer studies

In addition, our analysis also indicated that the hotspot mutations BRAF^V600K^, PIK3CA^N345K^, PIK3CA^E542K^, PIK3CA^E545K^, and PIK3CA^H1047L^ are promising candidates for neoantigens derived from cancer-relevant genes. Nearly half of all cutaneous melanomas carry activating BRAF^V600^ mutations, among which 10–30% contain the BRFA^V600K^ mutation, making it the second most common genotype after BRAF^V600E^ (33,34). BRAF^V600K^ lead to a gain in BraF protein function, as demonstrated by increased kinase activity, increased downstream signaling, and the ability to transform cells *in vitro* (35,36). Clinically, BRAF^V600K^ tumors cause patients to experience distant metastases sooner, and these patients have a higher risk of relapse and shorter survival than those with V600E tumors (37).

#### Identification of GIT cancer neoantigens

We further evaluated the genomic and transcriptomic data from colorectal cancer (CRC), esophageal squamous cell carcinoma (ESCC), and hepatocellular carcinoma (HCC) patients to further evaluate the putative neoantigens in these three types of cancers in the real world. We collected a total of 805 samples from different data sources (**Supplementary Table S6**). After quality checking, we analyzed a total of 691 samples, composed of 316 CRC samples, 290 ESCC samples, and 85 HCC samples. On average, we identified 17 (0, 316), 5 (0, 76), and 6 (0, 64) neoantigens by ImmuneMirror for each CRC, ESCC, and HCC patient, respectively. Noticeably, the neoantigen load was significantly correlated with favorable clinical outcome in terms of longer overall survival in ESCC samples (**Supplementary Figure S3**). CRC patients can be categorized as high microsatellite instability (MSI-H), low microsatellite instability (MSI-L) and microsatellite stability (MSS) according to the status of the mismatch repair pathway (38). MSI-H tumors respond well to immunotherapy, presumably due to a high TMB and neoantigen load (39,40). More interestingly, although the neoantigen load was not correlated with overall survival in CRC samples, we found that a subgroup of MSI-H CRC patients with MMR deficiency had a much lower neoantigen load for both HLA class I and II and a high TMB that was comparable to other MSI-H CRC patients (**Figure 4**). These patients were subject to advanced T stage (T4 vs. others: 30.8% vs. 0%, Fisher’s exact test *P* = 0.011).

**Figure 4.**
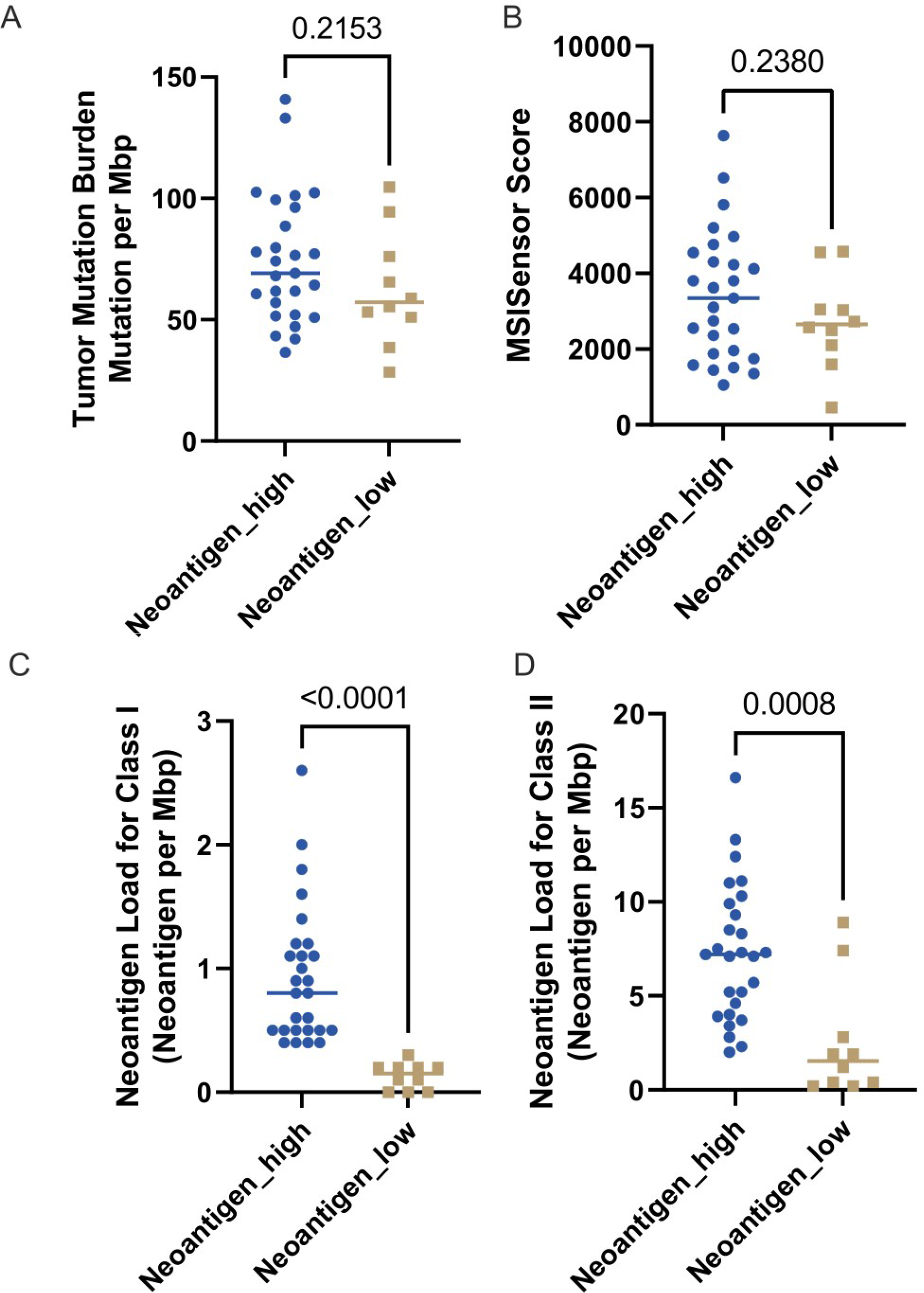
Lower neoantigen loads were detected in a subgroup of MSI-H CRC patients. A: TMB. B: MSISensor score for MSI status. C: The neoantigen load for HLA class I. D: The neoantigen load for HLA class II.

We identified a total of 12 putative neoepitopes that fulfilled Well’s criteria (8) and had an ImmuneMirror prediction score >0.5. These neoepitopes were derived from TP53, STAT3 and RAB35 with high affinity for the HLA-A*02:01, HLA-A*11:01, HLA-A*33:03, HLA-A*33:01, HLA-A*03:01, and HLA-A*02:06 HLA alleles (**Table 2**). More specifically, the neoepitope TP53^G245V^ (YMCNSSCMG<u>V</u>) restricted by HLA-A*02 was identified in the real-world data analysis of ESCC patient samples. This mutation affects the binding of p53 to DNA and interferes with the protein’s transcription activity. The RNASeq data indicated that this mutant is widely expressed in the tumor tissue (**Supplementary Figure S4**).

**Table 2:**
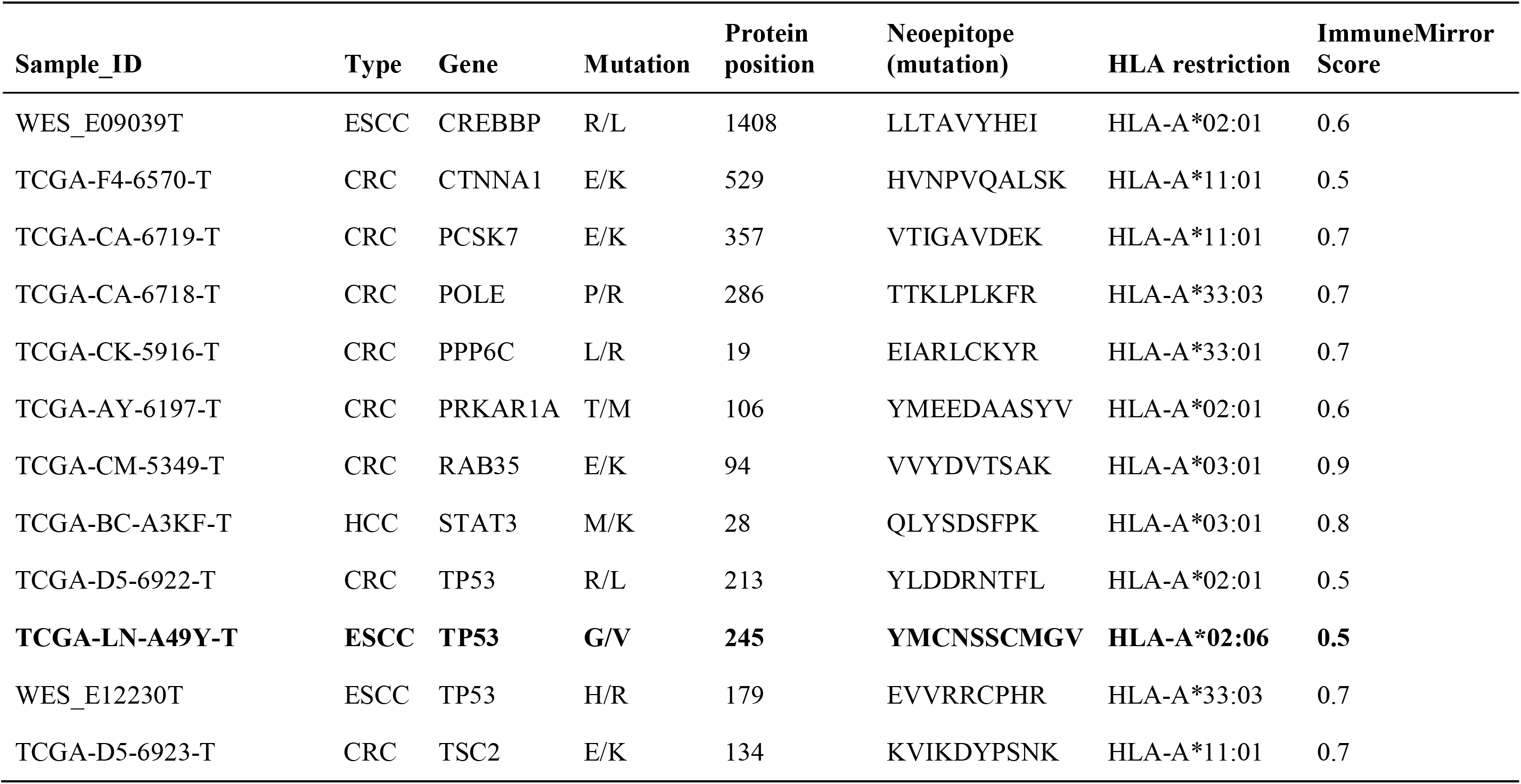
The neoantigens identified in GIT cancer samples

#### Validation of HLA-A02 binding with TP53.pG245V

We evaluated HLA-A02 binding affinity with neoepitopes derived from multiple mutations at G245 in TP53 using the QuickSwitch Quant HLA-A*02:01 Tretramer Kit-PE. The neoepitope TP53^G245V^ (YMCNSSCMG<u>V</u>) had a higher reference peptide exchange rate of 97.03% than the wild-type peptide YMCNSSCMGG (80.8%) (**Figure 5A**). The binding affinity of neoepitope-TP53^G245V^ (YMCNSSCMG<u>V</u>) was the highest among TP53^G245R^, TP53^G245D^, TP53^G245C^, and TP53^G245S^ (**Figure 5B**), and the Pearson’s correlation between the ImmuneMirror prediction scores and binding affinities was as high as 0.875 (**Figure 5C**). This result confirmed the effectiveness and reliability of ImmuneMirror as an advanced tool for neoantigen prediction.

**Figure 5.**
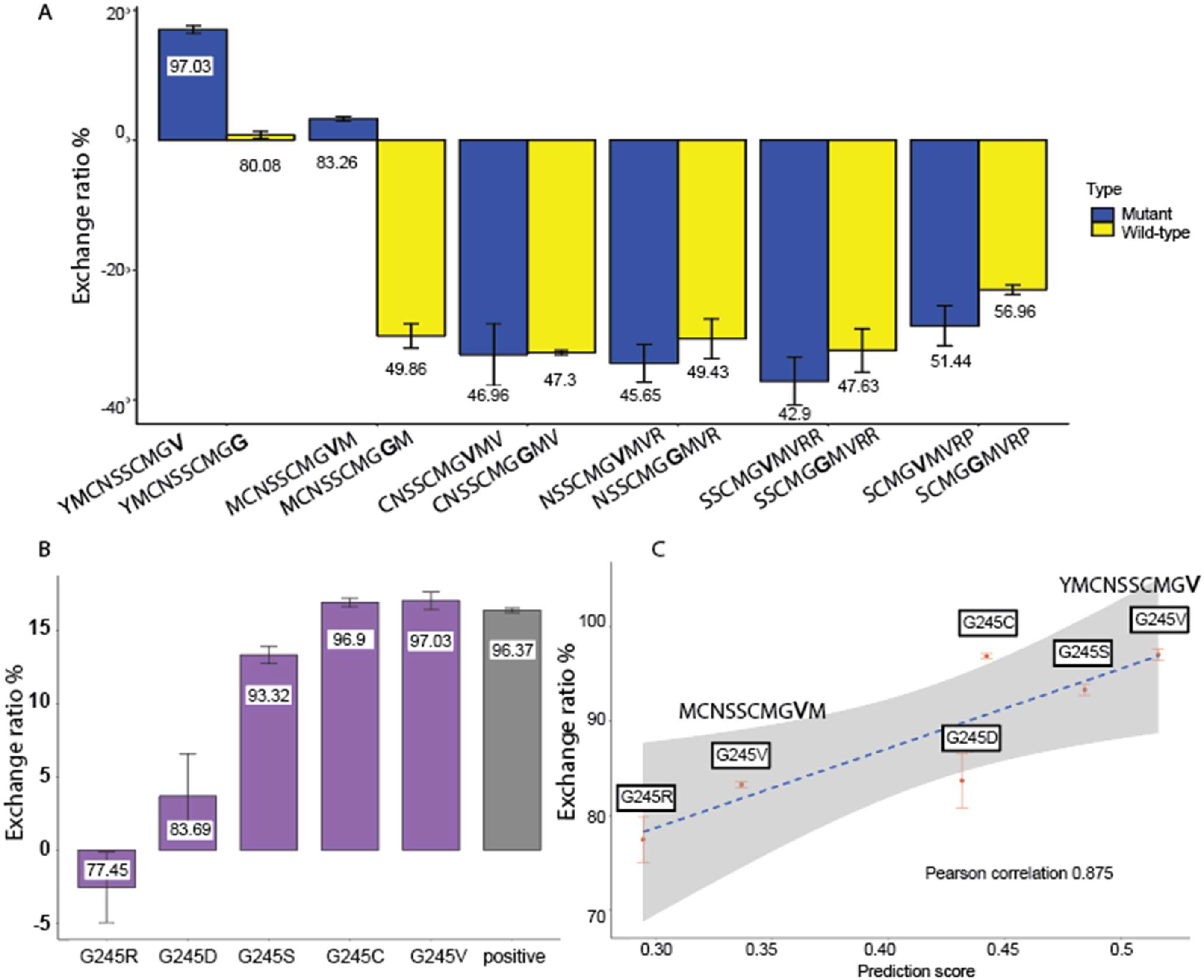
Validation of binding affinity between HLA-A02 and neoepitopes derived from TP53 mutations. **A)**The exchange ratios (mean ± SD) of the TP53^G245V^ mutant compared with the matched wild type (exchange ratio over 80% is used as the cutoff for positive and negative values). **B)**The exchange ratio (mean ± SD) of TP53^G245^ mutants compared with the positive control (an exchange ratio of 80% was considered biologically relevant). **C)**Scatterplot of the prediction score versus the exchange ratio (mean ± SD) for TP53^G245^ mutants.

## DISCUSSION

By integrating a balanced random forest model, we developed ImmuneMirror as a self-standing open-source pipeline and a web server for neoantigen prediction and prioritization. ImmuneMirror was trained and tested using immunogenic neoantigens collected from 19 studies. To our knowledge, this is the largest study to comprehensively evaluate the neoantigen prediction model using experimentally validated neopeptides to date. Accurate neoantigen prediction depends on including the biological features that essentially govern epitope immunogenicity. Referring to published studies, our model thoroughly integrates important biological features of immunogenic neoantigens. Additionally, we applied statistical methods to keep the most relevant features and developed a prediction model based on advanced machine learning algorithms. The effectiveness and reliability of ImmuneMirror have been confirmed by analyzing 805 samples of gastrointestinal tract cancers and experimental validation of selected candidates.

Both the ImmuneMirror and Wells’ study (8) models indicate that neopeptides with strong MHC binding affinity, long half-life and low agretopicity are most likely to be neoantigens. However, when analyzing the same data set, Wells’ criteria (8) tend to be very stringent about binding affinity, agretopicity and peptide stability to accommodate the needs of high specificity for clinical application, while the ImmuneMirror model offers an integrated approach considering more relevant biological features, including binding affinity, ‘agretopicity’ (9,11) and ‘foreignness’ (12–14), hydrophobicity, binding stability, peptide processing, and transportation scores without cutoffs, which provides more potential candidates for the purpose of research for further downstream experimental validation.

In our real-world data analysis, we found that neoantigen load was a predictor of good clinical outcomes in ESCC patients. Although it is known that MSI-H is an important molecular biomarker for selecting CRC patients who may benefit from anti-PD-1/PDL-1 therapy, we further identified a subgroup of MSI-H CRC patients enriched for advanced T stage that had relatively low neoantigen loads for HLA class I and II by ImmuneMirror. Promising results for immunotherapy have been demonstrated in a previous study that evaluated the efficacy of PD-1 blockade in advanced MSI-H patients across twelve different cancer types with an objective response rate in 53% of patients and complete response in 21% of patients; nevertheless, almost half of MSI-H cancer patients do not respond well to this treatment. This previous study also showed *in vivo* expansion of T-cell clones specifically activated by neoantigens in patient responses (40). Our results suggest that further stratification of MSI-H cancer patients based on neoantigen loads may be necessary, and a more detailed evaluation of the objective response rate of this unique subset of MSI-H patients to anti-PD-1/PDL-1 therapy is needed in a clinical trial.

The TP53^G245V^ mutation occurs at a total frequency of 0.13% in diverse cancers, such as diffuse glioma, non-small cell lung cancer, bladder urothelial carcinoma, endometrial carcinoma, head and neck squamous cell carcinoma, pancreatic adenocarcinoma, and esophageal squamous cell carcinoma, according to the records in the cBio Cancer Genomic portal (41). The discovery of the neoepitope TP53^G245V^ (YMCNSSCMG<u>V</u>) derived from this mutation restricted by HLA-A*02, a common HLA class I type in Caucasians and Asians, showed the effectiveness and great potential of ImmuneMirror for detecting the neoantigens. In addition to developing the neoantigen vaccine targeting this neoepitope, further identification of T cells that are specifically reactivated by this neoepitope is necessary for developing adoptive T-cell therapies for cancer patients carrying this specific mutation.

In summary, ImmuneMirror is an integrative analysis pipeline that can be applied for genomic and transcriptomic data analysis, especially for neoantigen prediction, in samples from a variety of cancer types. This tool could assist biologists in systematically evaluating the genomic and transcriptomic features relevant to the response to immunotherapy, including TMB, neoantigen load, MSI status, HLA typing, and the expression of the IPRES. More importantly, ImmuneMirror will be strategically useful as a guide for clinicians to tailor treatment strategies according to the genomic and transcriptomic profiles for precision medicine and to facilitate clinical trial design and patient selection with broad prospects for clinical applications. Additional experimental and clinical validation of the putative neoantigens identified in this study are warranted to determine the usefulness of these putative neoantigens for immunotherapy.

## DATA AVAILABILITY

The published article includes all data sets generated or analyzed during this study.

## Accession codes

### WES data

European Genome-phenome Archive (EGA): EGAS00001000932 (42); NCBI Sequence Read Archive (SRA): SRP033394 (43), NCBI Bioproject: PRJNA399748 (44); and TCGA ESCC, CRC and HCC samples from NCI Genomic Data Commons (https://portal.gdc.cancer.gov/). Data from a previous study carried out by Dai *et al*. (45).

### RNASeq data

TCGA ESCC, CRC and HCC samples from the NCI Genomic Data Commons (https://portal.gdc.cancer.gov/)

The open source ImmumeMirror pipeline and usage guide are available at https://github.com/weidai2/ImmuneMirror. The source code is released under the GNU General Public License version 3 (GPL >=3). The web server is freely available at http://immunemirror.hku.hk/App/ and does not have a login requirement.

## SUPPLEMENTARY DATA

Supplementary Data are available online.

## AUTHOR CONTRIBUTIONS

WD and ZL designed and supervised the study. GSC developed the ImmuneMirror pipeline, built the docker image, performed data analysis, tested the pipeline and web server, and compared ImmuneMirror with other similar pipelines. YG developed the machine learning model and performed the neoantigen analysis for frequent mutations derived from cancer-related genes. GSC and YG designed and developed the ImmuneMirror web server. GSC, YG, WD, and ZL wrote the manuscript. CLC and KOL are clinicians who advised on the analysis of the clinical specimens. NWK advised on experimental validation for the MHC class I binding assay. All the authors have read and approved the manuscript.

## Supplementary Figures

**Supplementary Figure S1**. ImmuneMirror pipeline workflow. The diagram focuses on the bioinformatics analysis steps and their workflows with intermediate processes involved, from raw FASTQ file preprocessing to neoantigen prediction and prioritization. RNA, Ribonucleic acid; WES, whole-exome sequencing; N, normal; T, tumor; L, lymph node; BAM, binary alignment map; QC, quality control; TPM, transcripts per million; SNV, single nucleotide variant; VCF, variant call format; Indel, insertion or deletion; MSI, microsatellite instability; HLA, human leukocyte antigen; MHC, major histocompatibility complex.

**Supplementary Figure S2**. Graphical analysis report of a patient sample produced by the ImmuneMirror pipeline as an example.

**Supplementary Figure S3**. Survival analysis of Pan-Cancer studies, including (A) CRC, (B) ESCC, and (C) HCC patients.

**Supplementary Figure S4**. The expression of the TP53^G245V^ mutation was detected by RNASeq as illustrated by Integrative Genomics Viewer (IGV).

## Supplementary Tables

**Supplementary Table S1**. The list of published studies used for construction and evaluation of the machine learning models for neoantigen prediction.

**Supplementary Table S2**. The list of peptides that were used for model training and testing.

**Supplementary Table S3**. The list of software and packages that were used in ImmuneMirror implementation.

**Supplementary Table S4**. Compatibility of ImmuneMirror with various operating systems and browsers.

**Supplementary Table S5**. Comparison of bioinformatics tools available for neoantigen prediction.

**Supplementary Table S6**. List of data sources for the WES and RNASeq data for this study.

