## Supplementary figures and images for "ImmuneMirror: a Machine Learning-based Integrative Pipeline and Web Server for Neoantigen Prediction"

### Supplementary Figure S1

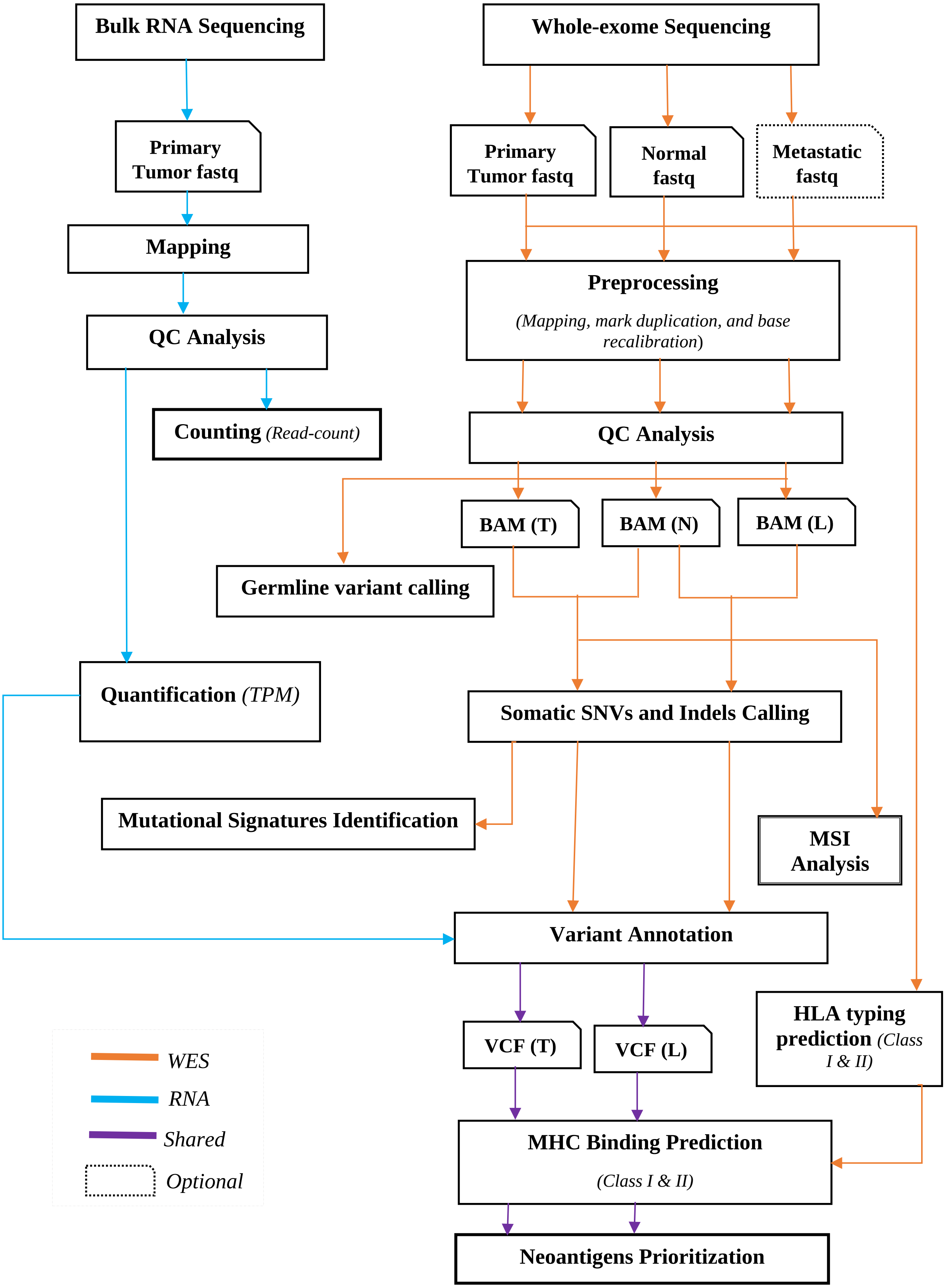

### Supplementary Figure S3

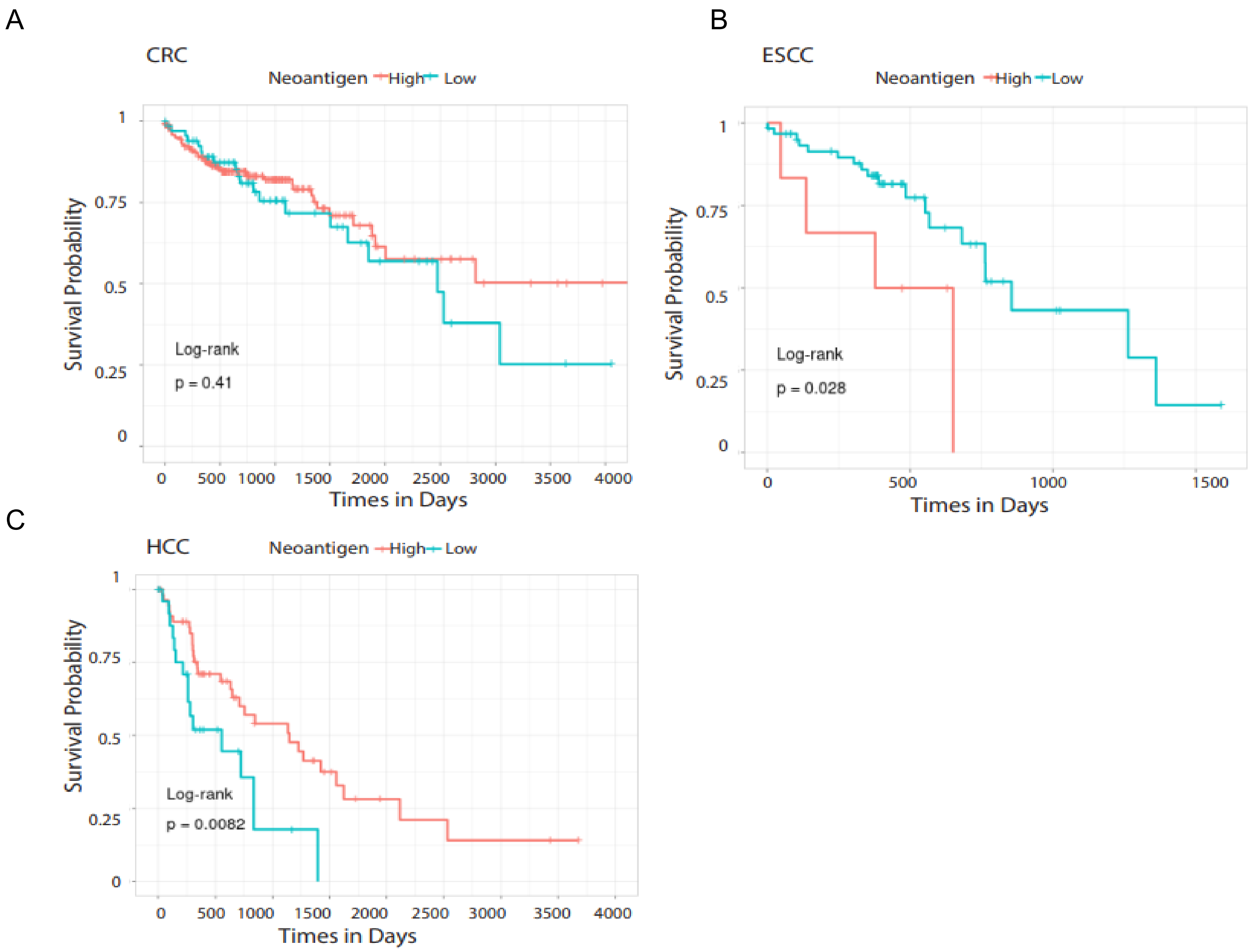

### Supplementary Figure S4

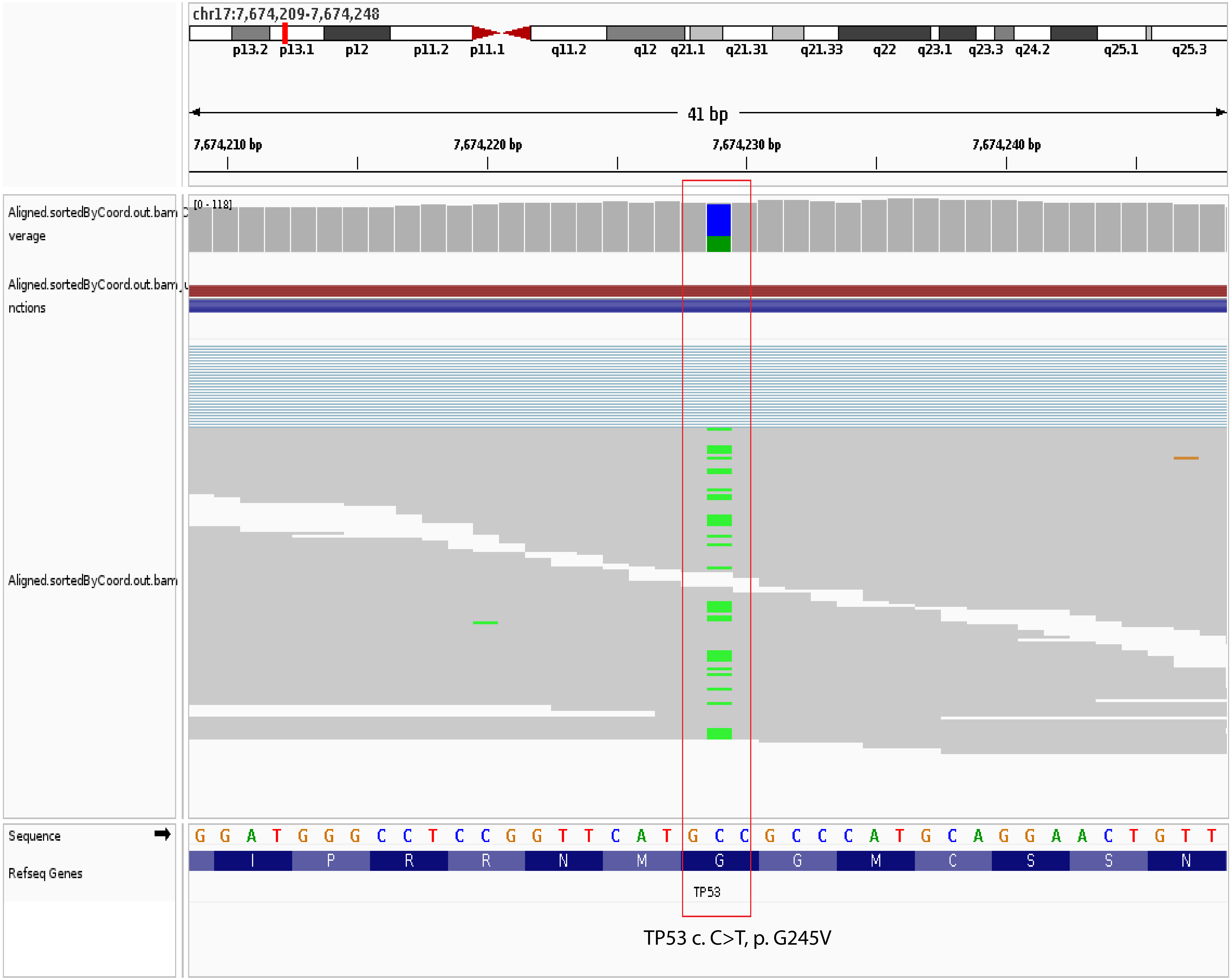


TP53 c. C>T, p. G245V
