## Supplementary Figure S2 for "ImmuneMirror: a Machine Learning-based Integrative Pipeline and Web Server for Neoantigen Prediction"

### Analysis Report

**Sample name: TCGA−AZ−4615−T**

##### A. Mutation Load

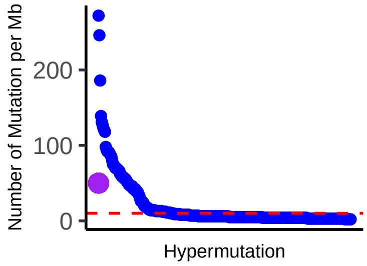


##### B. HLA types

| HLA Type I | HLA Type II |
|---|---|
| HLA−A*01:01 | DQA1*01:01 |
| HLA−A*24:02 | DQA1*05:01 |
| HLA−B*08:01 | DQB1*02:01 |
| HLA−B*44:05 | DQB1*05:01 |
| HLA−C*02:02 | DRB1*01:01 |
| HLA−C*07:01 | DRB1*03:01 |

##### C. Tumor neoantigen load for class I (without filtering)

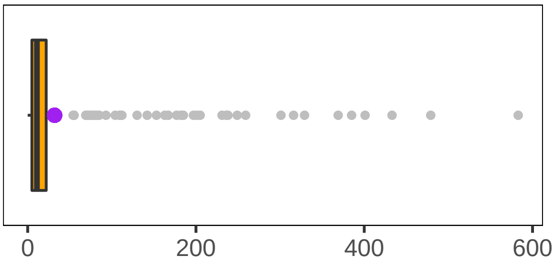


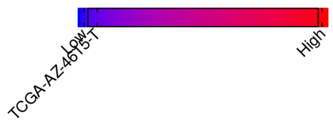


##### D. Tumor neoantigen load for class I (filtered)

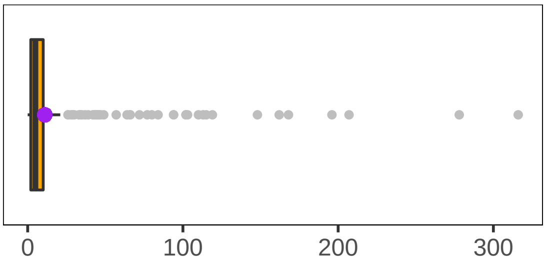


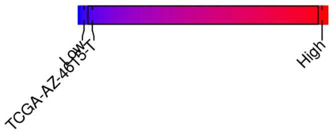


##### E. Tumor neoantigen load for class II (without filtering)

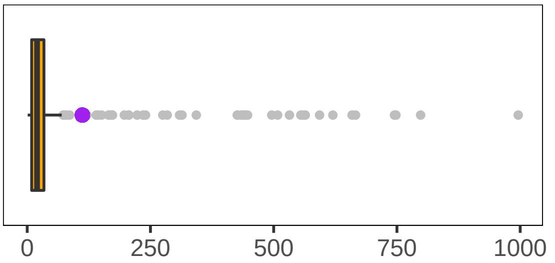


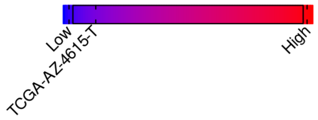


##### F. MMR status

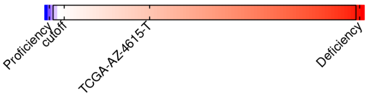


##### G. Germline mutation

| Gene | Mutation | AA_change | ClinVar |
|---|---|---|---|
| BRCA2 | − | − | − |
| B2M | − | − | − |
| JAK1 | − | − | − |
| JAK2 | − | − | − |
| PTEN | − | − | − |
| AKT1 | − | − | − |
| EGFR | c.C2011T | p.R671C | − |

##### H. Somatic mutation

| Gene | Mutation | AA_change |
|---|---|---|
| MLH1 | − | − |
| MLH3 | − | − |
| MSH6 | − | − |
| PMS2 | − | − |
| MSH2 | c.690T>C | p.C230C |

I. IPRES gene signature

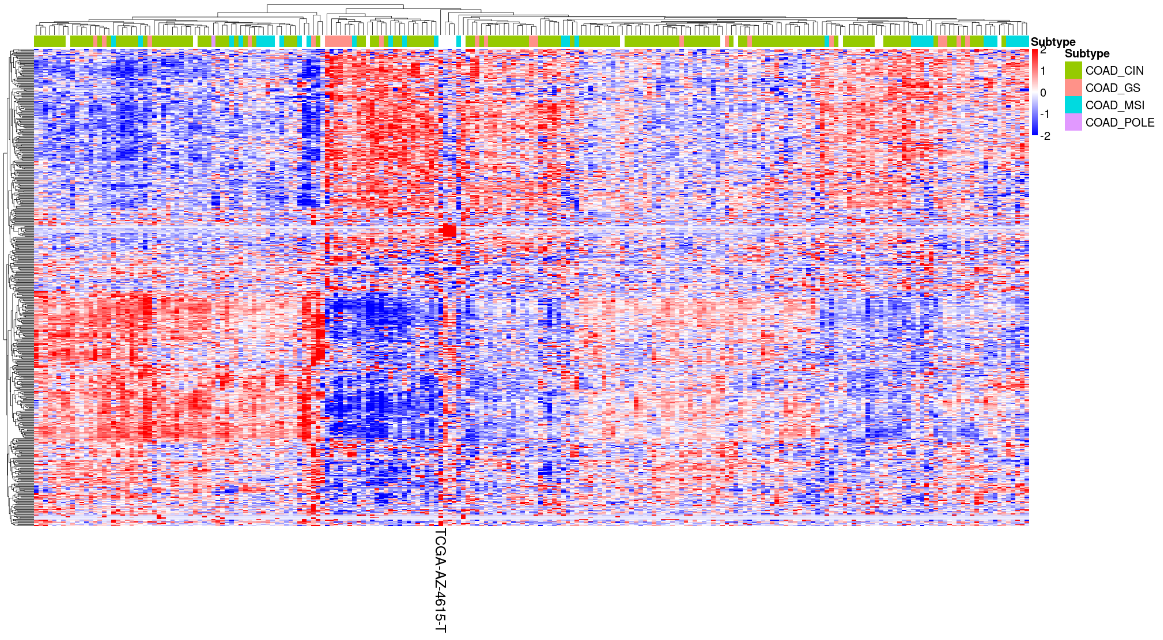


**Figure: Analysis report. Your sample is shown by purple dots. A. Mutation load. B. List of HLA types of your sample. C. Tumor neoantigen load for class I (without filtering). D. Tumor neoantigen load for class I (filtered). E. Tumor neoantigen load for class II (without filtering). F. MMR status. G. Germline mutation. H. Somatic mutation. I. IPRES gene signature.**

*Developed by*
*Dr. Wei Dai's group* (http://weidai-lab.hku.hk/)
Department of Clinical Oncology, The University of Hong Kong.
We acknowledge the *Health and Medical Research Fund (HMRF)*
(https://rfs1.fhb.gov.hk/english/funds/funds_hmrf/funds_hmrf_abt/funds_hmrf_abt.html) for support.
The work is jointly done with *Dr. Zhonghua Liu's group* (https://sites.google.com/view/drliu/home)
Department of Biostatistics, Columbia University, New York, NY, USA
