## Supplementary Data File for "ImmuneMirror: a Machine Learning-based Integrative Pipeline and Web Server for Neoantigen Prediction"

**Supplementary Material**

Detailed description of ImmuneMirror pipeline and Web server development

**Whole-exome sequencing**

***Preprocessing****:* The BWA-mem (1) aligner was used for short sequence reads mapping to the reference genome GRCh38 (hg38, <https://genome.ucsc.edu/>). The mapped reads were sorted and indexed using samtools (2) sort and index command, respectively. Next, the Picard Toolkit (http://broadinstitute.github.io/picard/) was used to mark the duplicated reads from sorted-indexed BAM files. To recalibrate base quality scores, we used the Genome Analysis Toolkit (GATK4) (3) in the final step of preprocessing module.

***Quality control of the aligned reads****:* We used the CollectHsMetrics command in Picard Toolkit to produce the hybrid-selection (HS) metrics from aligned BAM files. To predict the high quality neoantigens, the BAM files were discarded those have average coverage (MEAN_BAIT_COVERAGE) less than 30 and 40 for normal and tumor samples, respectively.

***Microsatellite instability analysis***: For microsatellite instability (MSI) analysis, the MSIsensor-pro (4) was applied to mapped tumor-normal pair samples.

***Germline variant calling***: We followed the GATK best practice workflow for the germline variant calling analysis using the human reference genome (GRCh38). The GATK4 (3) HaplotypeCaller command was used for germline SNPs (single-nucleotide polymorphism) and INDELs (frameshift insertions and deletions) calling in genomic variant call format (GVCF), and, next, the GATK GenotypeGVCFs tool was queried to perform the genotyping assembly in VCF file format from the pre-calculated GVCF file.

***HLA typing prediction*:**We applied the OptiType (5) and PHLAT (6) algorithms to determine HLA class I and class II haplotyping predictions, respectively, to each of the FASTQ files.

***Calling Somatic SNVs and Indels***: we used Mutect2 (GATK4) (3) for somatic SNVs and Indels calling in both the cases, i.e., tumor with matched normal and tumor with metastatic. The FilterMutectCalls (3) was applied for filtering the raw output form Mutect2. Moreover, the *SelectVariants* command in GTAK4 was performed to select the biallelic sites from filtered VCF output files, and the variants passed quality filtering were selected by VCFtools (7). For functional annotation analysis, the Funcotator (GATK4) was applied to allelic files that produced files in Mutation Annotation Format (MAF) as outputs.

***Variant annotation analysis***: The biallelic pass VCF files were annotated with RNA-seq TPM file using the VEP (8) tool.

***MHC binding prediction***: prior predicted HLA class I and class II alleles were utilized for corresponding MHC complexes predicting using pVACseq tool as implemented in pVACtools (9). We used the NetMHCpan, NetMHC, NetMHCcons, PickPocket, SMM, SMMPMBEC, MHCflurry, MHCnuggest methods for MHC class I binding predictions and the algorithms NetMHCIIpan, SMMalign, NNalign, and MHCnuggetsII for predicting the binding in MHC class II complexes as implemented in pVACseq. Next, The NetChop (10) tool was used for predicting the cleavage sites and TAP transportation in MHC class I predicted results. Finally, we implemented a machine learning model as a R function for prioritizing the neoantigens for HLA class I.

**RNA sequencing**

***Alignment***: The STAR aligner (11) was used for mapping RNA-seq FASTQ files with reference transcriptomes sequence (12).

***Quality Control*:** The CollectRnaSeqMetrics and CollectAlignmentSummaryMetrics commands in Picard Toolkit (<http://broadinstitute.github.io/picard/>) were queried in mapped BAM files for producing RNA alignment metrics and summary of alignment metrics, respectively.

***Quantification*:** The RNA-seq raw reads FASTQ files were quantified using *Salmon* (13) tool that can produce gene-level expression as TPM file. We also queried the HTSeq (14) tool to the BAM files for generating the gene-level expression in count-table format.

**Web server interface design**

The ImmuneMirror web server was implemented using Shiny web framework (<https://CRAN.R-project.org/package=shiny>). Various R packages were used for designing the interactive web interface, such as *shinyjs* (<https://CRAN.R-project.org/package=shinyjs>), *shinythemes* (<https://CRAN.R-project.org/package=shinythemes>), and *shinyFeedback* (<https://CRAN.R-project.org/package=shinyFeedback>). The R package *emayili* (https://datawookie.github.io/emayili/) was used for sending automatic email from the server to users.
