## Supplementary Table S1 for "ImmuneMirror: a Machine Learning-based Integrative Pipeline and Web Server for Neoantigen Prediction"

| **Index** | **Publication date** | **Reference** | **Author** | **Journal** | **Set** |
| --- | --- | --- | --- | --- | --- |
| 1. | 2013–05 | (1) | Robbins et al. | Nat Med | training set |
| 2. | 2013–11 | (2) | Van Roij et al. | J Clin Oncol |  |
| 3. | 2014–03 | (3) | Wick et al. | Clin Cancer Res |  |
| 4. | 2014–06 | (4) | Rajasagi et al. | Blood |  |
| 5. | 2014–07 | (5) | Lu et al. | Clin Cancer Res |  |
| 6. | 2015–04 | (6) | Rizvi et al. | Science |  |
| 7. | 2015–10 | (7) | Cohen et al. | J Clin Invest |  |
| 8. | 2016–01 | (8) | Kalaora et al. | Oncotarget |  |
| 9. | 2016–05 | (9) | Strønen et al. | Science |  |
| 10. | 2016–05 | (10) | Bassani-Sternberg et al. | Nature Commun |  |
| 11. | 2017–07 | (11) | Ott et al. | Nature |  |
| 12. | 2016–11 | (12) | Gros et al. | Nature medicine |  |
| 13. | 2017–07 | (13) | Le et al. | Science |  |
| 14. | 2015-05 | (14) | Carreno et al. | Science |  |
| 15. | 2020-11 | (15) | Cafri et al. | J Clin Invest |  |
| 16. | 2018-08 | (16) | Danilova et al. | Cancer Immunol Res |  |
| 17. | 2019-11 | (17) | Sneddon et al. | Oncoimmunology. |  |
| 18. | 2020-10 | (18) | Wells et al. | Cell | testing set |
| 19. | 2019-11 | (19) | Linette et al. | PNAS |  |

**Supplementary Table S1**: The list of published studies used for construction and evaluation of the machine learning model for neoantigen prediction.

**References**
