## Supplementary Table S3 for "ImmuneMirror: a Machine Learning-based Integrative Pipeline and Web Server for Neoantigen Prediction"

**Supplementary Table S3**. The list of software and packages were used in ImmuneMirror implementation.

| **Software and Algorithms** | **Source** | **Identifier** |
| --- | --- | --- |
| Docker version 19.03.8 | Docker, Inc. (Boettiger, 2015; Merkel, 2014) | http://www.docker.com |
| GNU bash version 5.0.17(1) | Free Software Foundation, Inc. | https://www.gnu.org/software/bash/ |
| Burrows-Wheeler Aligner (BWA) version 0.7.17 | (Li and Durbin, 2009) | http://bio-bwa.sourceforge.net/bwa.shtml |
| SAMtools version 1.10 | (Danecek et al., 2021) | https://github.com/samtools/samtools |
| Picard tools version 2.17.4 | Broad Institute | https://broadinstitute.github.io/picard/ |
| GATK (GenomeAnalysisTK) version 4.1.8.0 | Broad Institute (McKenna et al., 2010) | https://gatk.broadinstitute.org/hc/en-us |
| ANNOVAR | (Wang et al., 2010) | https://annovar.openbioinformatics.org/ |
| Python version 3.7.6 | Python Software Foundation | https://www.python.org/ |
| R version 3.6.3 | R Core Team (Ihaka and Gentleman, 1996) | https://www.r-project.org/ |
| razers3 version 3.5.8 | (Weese et al., 2012) | http://www.seqan.de/razers/ |
| OptiType 1.3.3 | (Szolek et al., 2014) | https://github.com/FRED-2/OptiType |
| PHLAT version 1.0 | (Bai et al., 2018) | https://sites.google.com/site/phlatfortype/ |
| JAVA version 1.8.0_275 | OpenJDK Community | https://openjdk.java.net/ |
| VCFtools version 0.1.16 | (Danecek et al., 2011) | https://github.com/vcftools/vcftools |
| HTSlib version 1.10.2 | (Bonfield et al., 2021) | https://github.com/samtools/htslib |
| ensembl-vep version 1.0 | (McLaren et al., 2016) | https://github.com/Ensembl/ensembl-vep |
| Perl version 5.30.0 | (Wall et al., 2000) | http://www.perl.org/ |
| MSIsensor-pro version 1.1 | (Jia et al., 2020) | https://github.com/xjtu-omics/msisensor-pro |
| pVACtools version 2.0.2 | (Hundal et al., 2020) | https://pvactools.readthedocs.io/en/latest/ |
| ncbi-blast version 2.11.0 | National Center for Biotechnology Information, U.S. National Library of Medicine | https://ftp.ncbi.nlm.nih.gov/blast/executables/blast+/LATEST/ |
| antigen.garnish version 2.0.0 | (Richman et al., 2019) | https://github.com/andrewrech/antigen.garnish |
| VAtools 4.1.0 | --- | <https://github.com/griffithlab/VAtools> |
| STAR version 2.7.5c | (Dobin et al., 2013) | https://github.com/alexdobin/STAR |
| salmon version 1.3.0 | (Patro et al., 2017) | https://combine-lab.github.io/salmon/ |
| HTSeq version 0.9.1 | (Anders et al., 2015) | <https://htseq.readthedocs.io/en/master/> |
| BCFtools version 1.10.2 | (Danecek et al., 2021) | https://github.com/samtools/bcftools |
| BEDtools version 2.29.2 | (Quinlan and Hall, 2010) | https://github.com/arq5x/bedtools2 |
| vcf2maf version 1.6.17 | (Kandoth, 2020) | https://github.com/mskcc/vcf2maf |
| NetChop version 3.0 | (Nielsen et al., 2005) | https://downloads.iedb.org/tools/netchop/3.0 |
