## Supplementary Table S4 for "ImmuneMirror: a Machine Learning-based Integrative Pipeline and Web Server for Neoantigen Prediction"

| **OS** | **Version** | **Chrome** | **Firefox** | **Microsoft Edge** | **Safari** |
| --- | --- | --- | --- | --- | --- |
| Linux | Ubuntu 20.04 | 106.0.5249.103 | 104.0 | n/a | n/a |
| macOS | HighSierra | [98.0.4758.109](http://98.0.4758.109/) | 104.0 | n/a | 12.0 |
| Windows | 10 | 106.0.5249.103 | 104.0 | 105.0.1343.53 | n/a |

**Supplementary Table S4**. Compatibility of ImmuneMirror with various operating systems and browsers.
