## Supplementary Table S5 for "ImmuneMirror: a Machine Learning-based Integrative Pipeline and Web Server for Neoantigen Prediction"

**Supplementary Table S5**. Comparison of bioinformatics tools available for neoantigen prediction.

| **Index** | **Software** | **Input** | **Raw data type** | **Source** | **Method for Prioritization** | **Docker**  **image** | **Web server/app** | **Class I prediction** | **Class II prediction** | **Multiple prediction methods** |
| --- | --- | --- | --- | --- | --- | --- | --- | --- | --- | --- |
| 1 | **ImmuneMirror** | Fastq, Vcf | WES, RNASeq | Mutation | **✓** | **✓** | **✓** | **✓** | **✓** | **✓** |
| 2 | **NeoPredPipe** | Vcf | -- | Mutation |  |  |  | **✓** | **✓** |  |
| 3 | **MuPeXI** | Vcf | -- | Mutation |  |  | **✓** | **✓** |  |  |
| 4 | **TSNAD** | Fastq | RNASeq | Gene fusion |  | **✓** |  | **✓** |  |  |
| 5 | **pVAC-Seq** | Vcf | -- | Mutation | **✓** | **✓** |  | **✓** | **✓** | **✓** |
| 6 | **CloudNeo** | Vcf, Bam | RNASeq | Mutation |  |  | **✓** | **✓** |  |  |
| 7 | **TIminer** | Vcf, Fastq | RNASeq | Mutation | **✓** |  |  | **✓** |  |  |
| 8 | **INTEGRATE-Neo** | Fastq, | WGS, RNASeq | Gene fusion |  |  |  | **✓** |  |  |
| 9 | **Neopepsee** | Fastq, vcf | RNASeq | Mutation | **✓** |  |  | **✓** |  |  |
| 10 | **Vaxrank** | Vcf, Bam | RNAseq | Mutation |  |  |  | **✓** |  |  |
| 11 | **OpenVax** | Fastq | WES, RNASeq | Mutation | **✓** | **✓** |  | **✓** | **✓** | **✓** |
| 12 | **TruNeo** | Fastq | WES, RNASeq | Mutation, gene fusion | **✓** |  |  | **✓** |  | **✓** |
| 13 | **ScanNeo** | Bam | RNASeq | Indels |  |  |  | **✓** |  | **✓** |
| 14 | **NeoFuse** | Fastq | RNASeq | Gene fusion |  |  |  | **✓** |  |  |
