## Supplementary Table S6 for "ImmuneMirror: a Machine Learning-based Integrative Pipeline and Web Server for Neoantigen Prediction"

| **Cancer type** | **No. of Patients** | **References** |
| --- | --- | --- |
| CRC | 359 | https://portal.gdc.cancer.gov/ |
| ESCC | 57 | (1) |
| ESCC | 19 | SRP033394 (2) |
| ESCC | 113 | EGAS00001000932 (3) |
| ESCC | 80 | https://portal.gdc.cancer.gov/ |
| ESCC | 78 | PRJNA399748 (4) |
| HCC | 99 | https://portal.gdc.cancer.gov/ |

**Supplementary Table S6**. The list of data sources for the WES and RNASeq data for this study.
